# A hierarchical Bayesian latent class mixture model with censorship for detection of linear temporal changes in antibiotic resistance

**DOI:** 10.1101/705897

**Authors:** Min Zhang, Chong Wang, Annette O’Connor

## Abstract

Identifying and controlling the emergence of antimicrobial resistance (AMR) is a high priority for researchers and public health officials. One critical component of this control effort is timely detection of emerging or increasing resistance using surveillance programs. Currently, detection of temporal changes in AMR relies mainly on analysis of the proportion of resistant isolates based on the dichotomization of minimum inhibitory concentration (MIC) values. In our work, we developed a hierarchical Bayesian latent class mixture model that incorporates a linear trend for the mean log_2_MIC of the non-resistant population. By introducing latent variables, our model addressed the challenges associated with the AMR MIC values, compensating for the censored nature of the MIC observations as well as the mixed components indicated by the censored MIC distributions. Inclusion of linear regression with time as a covariate in the hierarchical structure allowed modelling of the linear creep of the mean log_2_MIC in the non-resistant population. The hierarchical Bayesian model was accurate and robust as assessed in simulation studies. The proposed approach was illustrated using *Salmonella* enterica I,4,[5],12:i:- treated with chloramphenicol and ceftiofur in human and veterinary samples, revealing some significant linearly increasing patterns from the applications. Implementation of our approach to the analysis of an AMR MIC dataset would provide surveillance programs with a more complete picture of the changes in AMR over years by exploring the patterns of the mean resistance level in the non-resistant population. Our model could therefore serve as a timely indicator of a need for antibiotic intervention before an outbreak of resistance, highlighting the relevance of this work for public health. Currently, however, due to extreme right censoring on the MIC data, this approach has limited utility for tracking changes in the resistant population.

## Introduction

### Rationale

Identifying and controlling the emergence of antimicrobial resistance (AMR) is a high priority for researchers and public health officials. A critical component of this control effort is surveillance for emerging or increasing resistance, as evidenced by the number and scale of surveillance programs around the world [5] [6]. The aims of these surveillance programs are to enable detection of emerging resistance in a timely manner and to facilitate antimicrobial stewardship programs to be implemented properly and accurately [7]. Currently, detection of temporal changes in AMR relies primarily on analysis of the proportion of resistant isolates based on the dichotomization of minimum inhibitory concentration (MIC) [7]. The MIC can be obtained from laboratory methods or machine-learning approaches [7] [21]. Regardless of the approach to MIC determination or the breakpoint used, dichotomization results in a loss of information.

### Previous work and challenges

To date, the predominant approach to assessing changes in AMR has focused on evaluation of changes in the proportion of isolates resistant to a particular antibiotic over time. Several statistical methods have been employed, including the Cochran-Armitage trend test, logistic regression model with time as a co-variate [8] [9] [10], and the Mann-Kendall non-parametric method to test monotonic trends over time [11]. These statistical methods are based on MIC data that are dichotomized to resistant and non-resistant. Mazloom *et al.* [12] pointed out that methods based on categorizations cause information loss. As such, a focus on changes in proportion of bacteria that are categorized as resistance means that changes in the mean MIC of isolates above or below the resistant breakpoint, a phenomena referred to as MIC creep/decline, are not included in the current surveillance monitoring [13]. Similarly, reliance on dichotomized MIC data prevents monitoring of correlations in mean MIC, despite the fact that such information would aid in the identification of emerging joint resistance patterns.

Mean MIC estimation must address the natural characteristics of MIC values, which are obtained from serial dilution experiments or predicted using AI methods [21]. Regardless of the method for obtaining MIC, currently observed MIC values are interval censored. For example, an observed or predicted MIC of 8 for the organism A actually implies that the true MIC is > 4, ≤ 8, and ultimately unknown. Estimation of the mean MIC without adjusting for censorship is biased and likely to overestimate the bacterial resistance to an antibiotic [15]. An additional issue is modeling of the underlying distribution of the true unknown MIC values. With respect to MIC data, bacteria isolates typically consist of a mixture of two components. Depending upon the focus (or region), these two populations may be considered resistant and non-resistant populations or wild-type and non-wild-type populations; however, the presence of two overlapping populations is commonly considered reasonable for these bacteria. For each component, the true concentration of antibiotic required to inhibit bacterial growth (i.e., the true MIC value) is believed to follow a log-normal distribution [16]; hence, the log_2_MIC follows a normal distribution.

Statistical approaches for estimation of the mean MIC have been proposed previously. Kassteele *et al.* [17] suggested a model for mean log_2_MIC estimation that incorporated the censored nature of MIC data and adjusted for such bias using the interval-censored normal distribution as the underlying distribution. This model is a reasonable accommodation for censorship; however, the approach did not address the mixture of resistant and non-resistant populations in the observed data. Craig [18] proposed that the underlying distribution of log_2_MIC can be modeled by a mixture of Gaussian distributions with resistant and non-resistant populations. Jaspers *et al.* [1] [2] [3] modeled the full continuous MIC distribution for wild-type and non-wild-type bacteria populations determined by epidemiological cut-off rather than clinical breakpoints. According to their definition of bacterial categorization, the non-wild-type population has less informative distributions and was therefore estimated in non-parametric ways. These previously published approaches suggest that log_2_MIC mean can be estimated, although none of the above mentioned studies evaluated an approach for estimating a temporal trend in mean log_2_MIC, and such an approach is clearly a critical need for AMR surveillance programs. Therefore, building upon these previous studies, we sought to describe an approach to estimate the mean log_2_MIC and describe temporal changes in AMR, while simultaneously addressing the censored nature of MIC data and the mixed distributions of populations.

### Contribution

In this paper, we proposed a hierarchical Bayesian latent class mixture model with censoring to detect temporal changes in AMR. This proposed model was applied to data from human samples from the Center for Disease Control National Antimicrobial Resistance Monitoring System (NARMS) surveillance program and swine samples from the Iowa State University Veterinary Diagnostic Laboratory (ISU VDL). The human data consisted of *Salmonella* I,4,[5],12:i:- tested with ceftiofur (TIO) and chloramphenicol (CHL). The swine data included *Salmonella* I,4,[5],12:i:- tested with TIO. Our applications revealed some interesting patterns in the Results Section. Simulation studies showed that our model was accurate and robust in the estimation of the mean log_2_MIC in non-resistant populations, the linear temporal trend in non-resistant populations, and the proportion of resistant bacteria over time. Future work stemming from our model is also discussed in the Discussion Section. Inclusion of such analyses into current surveillance programs would provide additional insight for monitoring of temporal changes in AMR and would increase the value of information extracted from such surveillance systems.

## Methods

### Model notation and assumptions

Our hierarchical Bayesian model for detection of linear temporal changes in AMR must take into account the censored nature of the data and the underlying mixed distribution of the observations. The commonly used approach for analysis of two-fold serial dilution observations is to transform to base 2 logarithm. To account for censoring statistically, each observed MIC value was assumed to represent an interval of true MIC values rather than a single discrete point value. With the following notations, Table 1 explains the conversion between the observed log_2_MIC values and a continuous scale interval (*l*_*ij*_, *u*_*ij*_) for each isolate and each antibiotic.

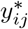: observed value of log_2_MIC for isolation *j* in year *i*;
*y*_*ij*_: latent true value of log_2_MIC for isolation *j* in year *i*;
*l*_*ij*_, *u*_*ij*_: lower bound and upper bound of the true value *y*_*ij*_, *y*_*ij*_ ∈ (*l*_*ij*_, *u*_*ij*_); and
*c*_*ij*_: latent indicator of the bacterial population from which the observation was drawn, where *i* = 1, 2, …*I*, and *j* = 1, 2, …, *n_i_*. Here, *I* is the total number of years, and *n*_*i*_ is the total number of isolates tested in the *i*th year.

**Table 1.** Conversion table between observed and latent log_2_MIC.

| $y_{ij}^*$ | censor type | $l_{ij}$ | $u_{ij}$ | $y_{ij} \in (l_{ij}, u_{ij})$ |
| --- | --- | --- | --- | --- |
| $\leq a$ | left censored | $-\infty$ | $a$ | $-\infty < y_{ij} \leq a$ |
| $= a$ | interval censored | $a - 1$ | $a$ | $a - 1 < y_{ij} \leq a$ |
| $> a$ | right censored | $a$ | ∞ | $a < y_{ij} < \infty$ |

Denote *S* as the conversion function between 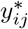 and *y*_*ij*_, and therefore, 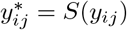, which is depicted in Fig 1 by the step-like plot. The distribution of the latent true value of log_2_MIC was assumed to be a bimodal Gaussian mixture model, which corresponds to the underlying mixture of resistant and non-resistant populations.

**Fig 1.**
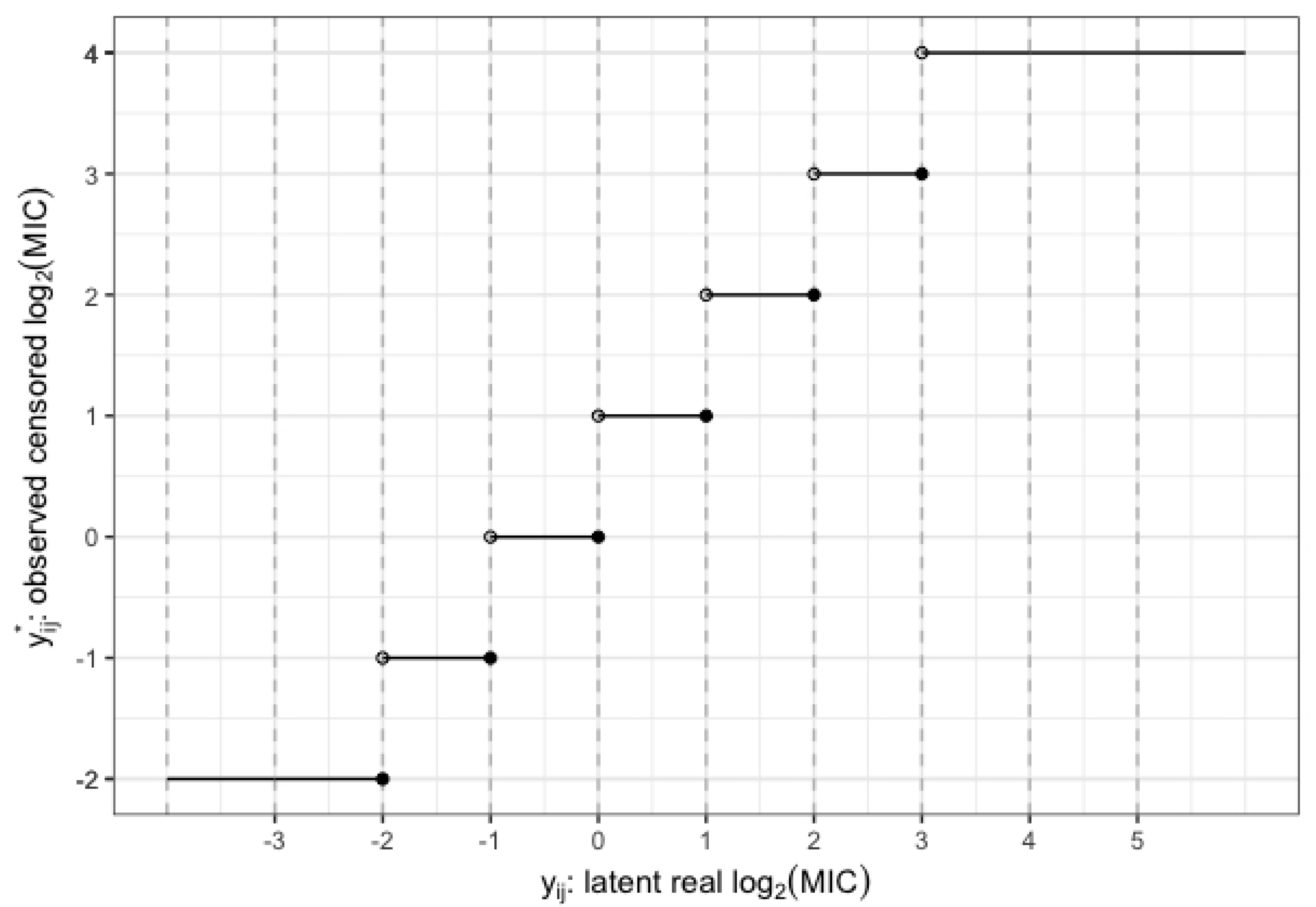
Conversion plot between observed and latent log_2_MIC. In this example, the serial dilution experiment starts at MIC = 2^−2^ = 0.25 and ends at MIC = 2^4^ = 16.

We proposed a model with a linear trend in the mean log_2_MIC over time. The motivation for this model stemmed from the results of a naïve analysis of *Salmonella* enterica I,4,[5],12:i:- and antibiotic CHL in the CDC NARMS dataset (the grey line displayed in Fig 2). Since the true log_2_MIC cannot be observed with the raw data due to censoring, the naïve mean log_2_MIC was calculated and presented over time. The naïve analysis for mean log_2_MIC ignored the nature of censoring of MIC data, resulting in calculation of the arithmetic average of log_2_MIC each year. For example, if the observed MIC value was ≤ 2, then log_2_MIC = log_2_(2) = 1 was treated as the corresponding log_2_MIC value and was therefore used for the naïve mean calculation. Although mathematically this approach has some issues, it served to illustrate the linear trend in the non-resistant population. The potential to observe and assess the presence of a linear trend is less useful in the resistant population, because the majority of the observed log_2_MIC results are right censored at the highest concentration of the serial dilutions. As a consequence, in the resistant population, the naïve mean log_2_MIC barely changed over time. Our assumption about the linear trend in the naïve mean log_2_MIC was only applicable to the non-resistant population.

**Fig 2.**
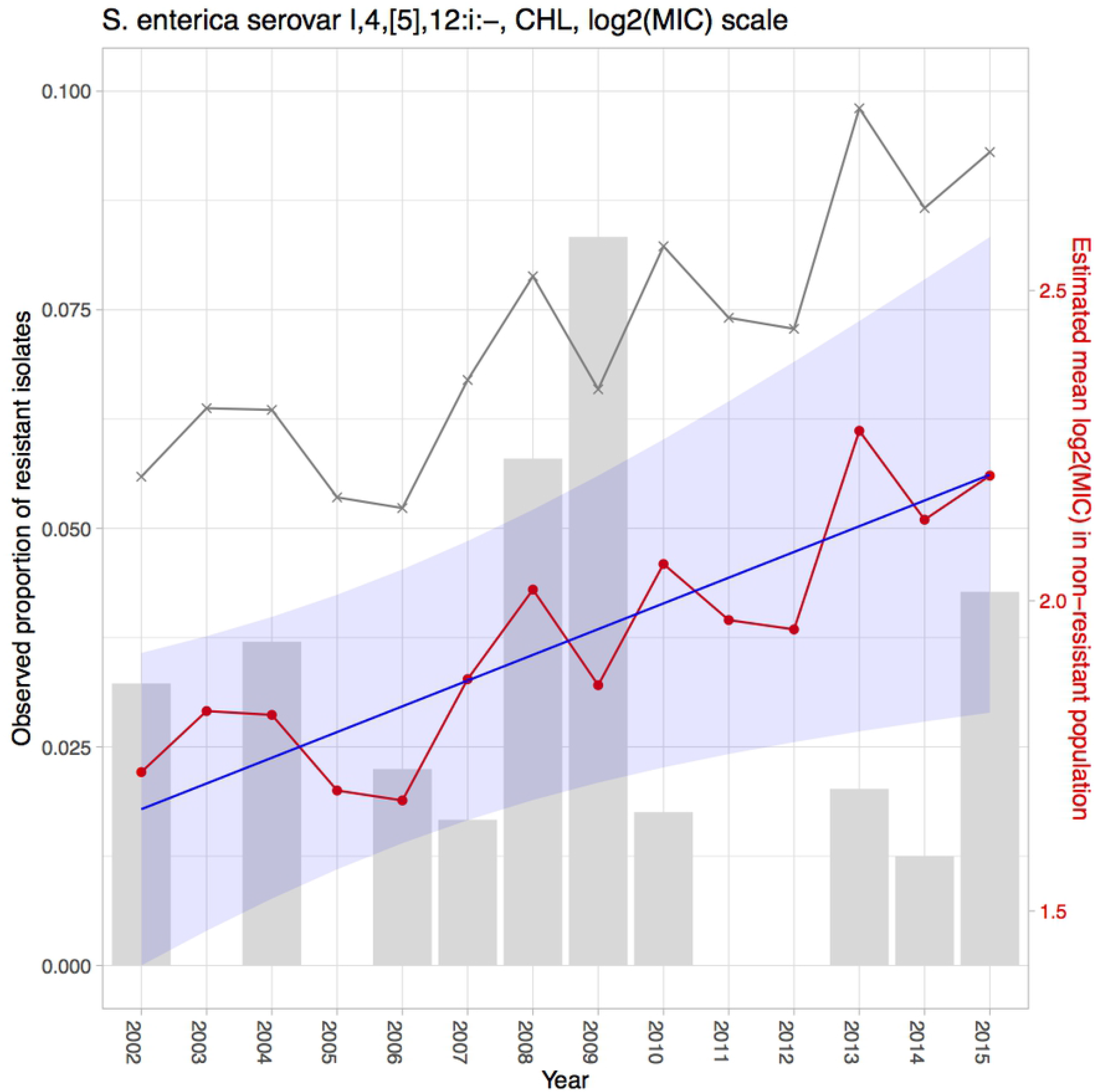
Estimation results for *Salmonella* enterica I,4,[5],12:i:- tested with CHL in the CDC NARMS dataset. The grey bars represent the observed proportions of resistant bacteria (left y-axis). The grey line indicates the naïve mean of log_2_MIC in the non-resistant population. The red line connects the point estimates of the mean log_2_MIC in the non-resistant population. The blue line represents the estimated linear trend, shaded with its 95% CI.

### Model description

Below, we describe in detail the approaches we used; however, as an introduction, we first provide a brief lay summary of the approach. In the first level of the model, we modeled the data with a mixture Gaussian distribution within each year. This level of model did not allow for estimation of a time effect, because each year was analyzed separately without being imposed by any pattern in time. The second level added complexity to the model via regression of the yearly mean log_2_MIC towards a line. For the resistant population, we assumed a flat line and that yearly randomness arose around this line. For the non-resistant population, we assumed a non-flat line that is a linear function of time. This time effect was estimated by a slope parameter in the model (described below): a positive result implies that the AMR is increasing with time, while a negative result implies that the AMR is decreasing with time. We used the real data for such a model with two levels and showed that the linear trend model was able to quantify the year effect observed in the year-to-year mode. Based on the notations and assumptions described in the beginning of the Methods Section, the construction procedure of the hierarchical Bayesian latent class mixture model with censoring and linear trends is described as follows:

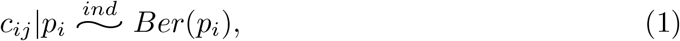

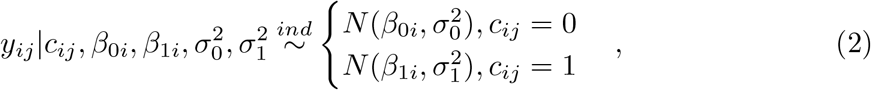

where *i* = 1, 2, …, *I*; *j* = 1, 2, …, *n_i_*. The variable before the pipe (“|”) was modeled with some distribution parameterized by the variable(s) behind the pipe. *Ber*(*p*) denotes a Bernoulli distribution with probability *p*, and *N* (*β*, *σ*^2^) denotes a normal distribution with mean *β* and variance *σ*^2^. In the *i*th year, the *j*th isolate comes from the resistant population with probability *p*_*i*_ and from the non-resistant population with probability 1 − *p_i_*. The parameters *β*_0*i*_ represent the mean log_2_MIC for the non-resistant group in *i*th year, and the parameters *β*_1*i*_ represent the mean log_2_MIC for the resistant group in the *i*th year. The variances for both components, 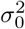 and 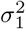, were set as invariant across the years, because we expected the spread of the observations within one population to be consistent over time. So far, the model allowed estimation of the mean log_2_MIC for each year but has not imposed any constraints on the yearly means. Considering the heterogeneity of bacteria isolates in the MIC dataset, due perhaps to different sampling collection methods from year to year or different labs used to test isolates (e.g., CDC NARMS dataset contains data collected from multiple institutes), a hierarchical modeling strategy was adopted to borrow information about mean log_2_MIC values across years and to integrate uncertainty from each individual year.

Based on the descriptive naïve means of log_2_MIC in the non-resistant population, we proposed to incorporate a linear trend into the model above to describe the temporal changes of the mean log_2_MIC in the non-resistant group for the organisms and antibiotics that appeared to be candidates for formal assessment of a linear pattern. First, we modeled the yearly mean log_2_MIC of the non-resistant population by introducing the hyper-parameters *γ*_0_ and *γ*_1_, with a simple linear model as follows:

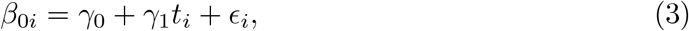

where 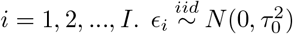. Time (year) was used as a covariate with *t*_*i*_ = *i*. For the first year of our observation *t*_*i*_ = *i* = 1. This is equivalent to

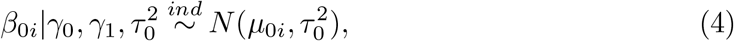

where *μ*_0*i*_ = *γ*_0_ + *γ*_1_*t_i_*. Second, we modeled the yearly mean log_2_MIC of the resistant population, using the hyper-parameter *μ*_1_ which is a constant:

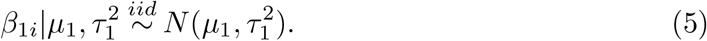

This model implied that the yearly mean of log_2_MIC in the non-resistant population is distributed around a straight line with intercept *γ*_0_ and slope *γ*_1_, with normally distributed error and variance quantified by 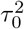, while the yearly mean of log_2_MIC in the resistant population is distributed around a constant *μ*_1_, also with normally distributed error and variance quantified by 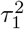. We modeled the yearly mean log_2_MIC of the resistant population with a constant instead of a linear trend because the MIC determination of the observed log_2_MIC is often highly right censored, meaning that we do not have enough information to estimate *β*_1*i*_ or its trend. If more studies reported end-point dilutions for resistant isolates, modeling the time trend in the resistant population would likely be feasible.

Further modeling of the first level, i.e. Eq (1) and Eq (2)), involved addition of more hyper-parameters in the hierarchical structure for the proportion of the resistant population in the *i*th year, *p_i_*. This parameter was modeled with a normal distribution through a logit link function:

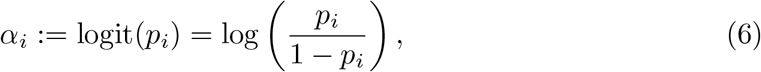

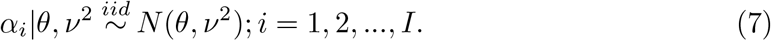

Let Θ be the vector of all unknown parameters 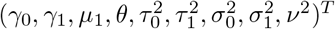; *f* be a generic expression for probability density function (pdf) or probability mass function (pmf). Also, 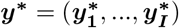 where 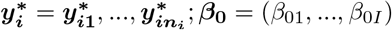 *β*_**1**_ = (*β*_11_, …, *β*_1*I*_); ***p*** = (*p*_0_, …, *p*_*I*_); *i* = 1, 2, …, *I*. The joint likelihood function was used as follows:

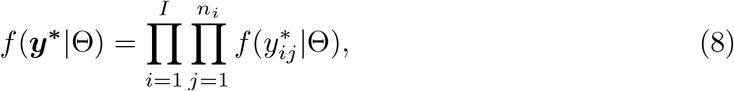

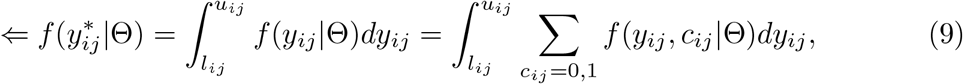

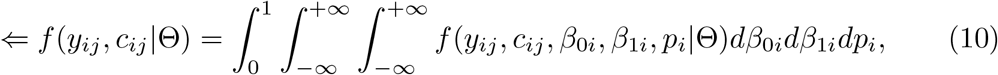

Based on densities and masses from Eq (1) to Eq (7):

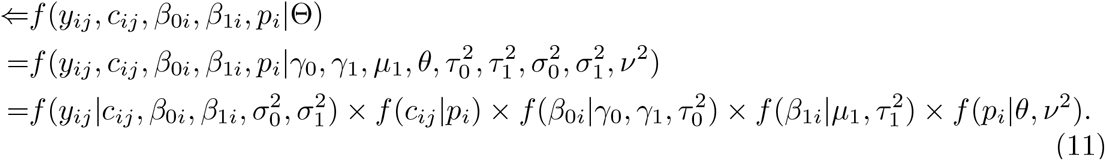

For Eq (9), our latent variables *y*_*ij*_ and *c*_*ij*_ were integrated (summed) over their possible range (values) to obtain the likelihood function of the observed data. In Eq (10), the mean and proportion parameters were also integrated over their supports. Eq (11) shows the derivation of the likelihood of latent variables and parameters from the data model.

### Prior distribution for hierarchical model parameters

The full Bayesian analysis required a joint prior distribution of all unknown parameters in the model. In our model setting, the vector of unknown parameters was 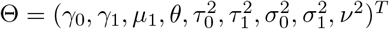. Furthermore, we assumed independent prior distribution for each parameter. The inverse gamma distribution was assigned to each variance of the data model and hierarchical part, due to their positive supports:

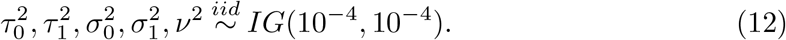

Non-informative prior was assigned to each of the linear parameters and mean parameters of the hierarchical part, because we did not have sufficient prior knowledge about these parameters. Their supports are described as follows:

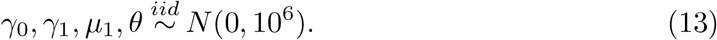

Using the Bayesian rule, our goal was to obtain samples and draw inference from the posterior distribution, which can be expressed based on densities from (8) to (13):

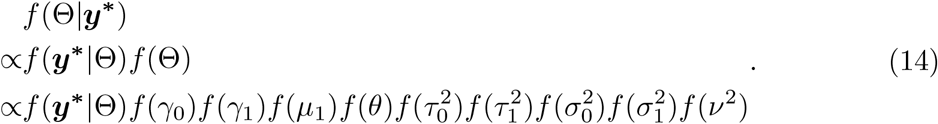

The posterior distribution did not have a closed form, and we illustrated the sampling approach in the following section with some real data applications.

## Application to real datasets

The goal of the assessment of the linear trend was to add a new dimension to understanding the temporal changes of AMR in both humans and livestock. To illustrate this, we applied the proposed model to human data from the CDC NARMS surveillance program and swine data submissions to the ISU VDL. The CDC NARMS data included *Salmonella* I,4,[5],12:i:- tested with CHL and TIO, while the swine data included *Salmonella* I,4,[5],12:i:- tested with TIO.

### Description of the CDC NARMS data used

NARMS program collects isolates of *Salmonella* spp., *Escherichia coli* and *Campylobacter* spp. The AMR data collected for the CDC surveillance program were obtained from bacteria isolated from patients who attend public health departments or hospitals that are part of the CDC NARMS surveillance network between 1996 and 2015. In the CDC NARMS dataset, the most abundant species was *Salmonella* enterica, which accounted for 58.70% of the 54351 total isolates. Serotype I,4,[5],12:i:- had 892 records in the dataset, accounting for 2.79% of the *Salmonella* enterica isolates. We chose *Salmonella* enterica I,4,[5],12:i:- because this strain is an emerging pathogen with public health implications for both hosts. The antibiotics CHL and TIO were selected for evaluation of a linear trend based on evidence of a linear trend in the naïve mean log_2_MIC in the non-resistant population descriptive analysis. For each of the two organism-antibiotic combinations, the distribution of the observed MIC was a mixture of two components: resistant and non-resistant populations. Therefore, the model assumptions were satisfied for these two combinations. Prior to the linear trend analysis, we discarded records before 2002, because the data were sparse for the first six years of the surveillance dataset (fewer than 10 isolates).

### Description of the VDL data used

Organisms isolated from livestock and submitted to veterinary diagnostic laboratories offer insight into the emergence and patterns of AMR. This population of organisms could offer different and unique information about AMR, providing insight into the implications for the spread of resistance through the local environment as well as occupational exposure. Yuan *et al.* (2018) reported that the swine population submitted to the ISU VDL suggested emergence of the pathogen *Salmonella* I,4,[5],i- sooner than the NARMS data. For comparison with the analysis for the CDC NARMS data and also for the sake of model assumptions, we applied the proposed model to a subset of ISU VDL data: *Salmonella* enterica I,4,[5],12:i:- tested with TIO from swine submissions. We did not evaluate CHL as for the CDC NARMS data, as this antibiotic was not used by the ISU VDL.

The ISU VDL dataset provided surveillance data on MIC from 2003 to 2018, comprising 93,634 isolates in total in the swine subset. Of these, 21,392 isolates are *Salmonella* enterica. Our target subset, *Salmonella* enterica serotype I,4,[5],12:i:- in the swine submissions included 1,967 isolates. We removed data obtained in 2018, because records in 2018 were not complete and still being processed in the diagnostic laboratory at the time of our analysis. We also excluded data from the first 8 y and focused on data obtained between 2011 and 2017, due to the sparsity of isolates observed in the first few years.

The implementation of the Bayesian model to the CDC NARMS dataset and the VDL dataset followed the same procedure. We describe the determination of the initial values of the Markov Chain Monte Carlo (MCMC) chain and calculation for inference in the next subsection.

### Implementation

Our proposed hierarchical Bayesian latent class mixture model with censoring and linear trend was implemented using the MCMC Gibbs sampling method. The Gibbs sampling algorithm was adapted for censorship in a finite mixture model [19] [20]. The algorithm of the Gibbs sampler is provided in S1 Appendix. All computation was implemented using R 3.3.5. All R scripts (including data cleaning, model construction, model implementation, results extraction, and results visualization) are provided on Github ^1^.

The initial values for MCMC were obtained from the raw data, so that the MCMC chain converged soon. For one combination of organism and antibiotic, the naïve mean of log_2_MIC in the non-resistant population in each tested year without censorship was calculated as the initial values for *β*_0*i*_. Similarly, the naïve mean of log_2_MIC values in the resistant population in each year was calculated as initial values for *β*_1*i*_, *i* = 1, 2,…, *I*. For the CDC NARMS dataset, *I* = 14, and for the VDL dataset, *I* = 7. The initial values for the linear parameters *γ*_0_ and *γ*_1_ were obtained from fitting *β*_0*i*_ and years *i* = 1, 2,…, *I* with simple linear regression. The estimated standard deviation of the error term of the simple linear regression was used as the initial value for *τ*_0_. The sample mean and sample standard deviation of *β*_1*i*_(*i* = 1, 2,…, *I*) were calculated as the initial values for *μ*_1_ and *τ*_1_, respectively. The initial values for the proportion of the resistant population *p*_*i*_ were calculated by dividing the number of resistant isolates by the total number of isolates in each year. The initial values of *α_i_*(*i* = 1, 2,…, *I*) were obtained by performing a logit transformation on the proportions *p_i_*.

Ten thousand iterations were performed, and the remaining 6,000 iterations after the 4,000 burn-in iterations were collected to make inferences. The parameters in the model were estimated by the mean of posterior distribution. The 2.5^th^ and 97.5^th^ percentiles of those 6,000 samples of posterior draws were used for determination of the 95% credible interval (CI). Table 2, Table 3, and Table 4 provide the point and interval estimates for the mean log_2_MIC for both resistant and non-resistant populations, the proportions of the resistant population; Table 5 provides the same set of information for the intercepts and slopes of the linear trends. These estimation results are also shown in Fig 2, Fig 3, and Fig 4 for better visualization.

**Table 2.** Point estimates and 95% CIs for mean and proportion parameters for *Salmonella* I,4,[5],12:i:- tested with (CHL) from the CDC NARMS dataset.

|  | Estimated mean log<sub>2</sub>MIC in the non-resistant population |  | Estimated mean log<sub>2</sub>MIC in the resistant population |  | Estimated proportion of resistant population |  |
| --- | --- | --- | --- | --- | --- | --- |
| Year | $\hat{\beta}_{0i}$ | 95% CI of $\hat{\beta}_{0i}$ | $\hat{\beta}_{1i}$ | 95% CI of $\hat{\beta}_{1i}$ | $\hat{p}_i$ | 95% CI of $\hat{p}_i$ |
| 2002 | 1.7238 | (1.5553, 1.8918) | 14.7834 | (7.9651, 22.0001) | 0.0263 | (0.0128, 0.0444) |
| 2003 | 1.8222 | (1.6545, 1.9887) | 14.6345 | (7.8851, 21.8576) | 0.0252 | (0.0084, 0.0384) |
| 2004 | 1.8161 | (1.6386, 1.9864) | 14.4699 | (8.4943, 21.7372) | 0.0265 | (0.0133, 0.0468) |
| 2005 | 1.6942 | (1.5195, 1.8615) | 14.6031 | (7.8794, 21.8553) | 0.0245 | (0.0067, 0.0333) |
| 2006 | 1.6782 | (1.5772, 1.7793) | 14.2337 | (8.6058, 22.2089) | 0.0252 | (0.0109, 0.0346) |
| 2007 | 1.8737 | (1.7584, 1.9877) | 14.6124 | (7.2965, 21.3269) | 0.0253 | (0.0096, 0.0370) |
| 2008 | 2.0180 | (1.9090, 2.1258) | 14.8148 | (8.0501, 22.2385) | 0.0287 | (0.0188, 0.0590) |
| 2009 | 1.8640 | (1.7291, 1.9975) | 14.6725 | (8.2464, 22.1023) | 0.0301 | (0.0210, 0.0711) |
| 2010 | 2.0592 | (1.9447, 2.1789) | 14.7990 | (8.4321, 21.7882) | 0.0253 | (0.0083, 0.0351) |
| 2011 | 1.9689 | (1.8522, 2.0900) | 14.5946 | (7.9317, 21.9562) | 0.0239 | (0.0061, 0.0308) |
| 2012 | 1.9539 | (1.8505, 2.0570) | 14.6309 | (7.9532, 22.0486) | 0.0239 | (0.0058, 0.0316) |
| 2013 | 2.2740 | (2.1836, 2.3674) | 14.8033 | (7.8064, 21.4324) | 0.0253 | (0.0128, 0.0336) |
| 2014 | 2.1307 | (2.0357, 2.2311) | 14.5674 | (8.3267, 21.7343) | 0.0248 | (0.0085, 0.0348) |
| 2015 | 2.2017 | (2.1175, 2.2860) | 14.4117 | (8.2791, 21.2827) | 0.0275 | (0.0185, 0.0478) |

**Table 3.** Point estimates and 95% CIs for mean and proportion parameters for *Salmonella* I,4,[5],12:i:- tested with (TIO) from the CDC NARMS dataset.

|  | Estimated mean log<sub>2</sub>MIC in the non-resistant population |  | Estimated mean log<sub>2</sub>MIC in the resistant population |  | Estimated proportion of resistant population |  |
| --- | --- | --- | --- | --- | --- | --- |
| Year | $\hat{\beta}_{0i}$ | 95% CI of $\hat{\beta}_{0i}$ | $\hat{\beta}_{1i}$ | 95% CI of $\hat{\beta}_{1i}$ | $\hat{p}_i$ | 95% CI of $\hat{p}_i$ |
| 2002 | -1.3309 | (-1.4990, -1.1657) | 12.7589 | (5.6945, 22.6436) | 0.0415 | (0.0246, 0.0596) |
| 2003 | -1.1844 | (-1.3495, -1.0210) | 12.7476 | (5.6395, 22.6380) | 0.0422 | (0.0258, 0.0624) |
| 2004 | -1.3890 | (-1.5780, -1.2017) | 12.5196 | (5.2118, 22.6231) | 0.0431 | (0.0271, 0.0654) |
| 2005 | -1.2564 | (-1.4195, -1.0979) | 12.7641 | (5.6486, 22.7206) | 0.0415 | (0.0256, 0.0583) |
| 2006 | -1.1858 | (-1.2808, -1.0922) | 12.8693 | (5.8527, 22.5644) | 0.0423 | (0.0287, 0.0586) |
| 2007 | -0.8702 | (-0.9801, -0.7550) | 12.7401 | (5.5777, 22.7752) | 0.0418 | (0.0242, 0.0603) |
| 2008 | -0.9226 | (-1.0280, -0.8147) | 12.7304 | (5.6264, 22.6573) | 0.0427 | (0.0274, 0.0622) |
| 2009 | -0.7899 | (-0.9158, -0.6651) | 12.7352 | (5.6360, 22.6448) | 0.0413 | (0.0246, 0.0588) |
| 2010 | -0.8039 | (-0.9231, -0.6844) | 12.7766 | (5.7197, 22.6692) | 0.0417 | (0.0250, 0.0576) |
| 2011 | -0.9334 | (-1.0518, -0.8116) | 12.7302 | (5.5366, 22.6482) | 0.0432 | (0.0269, 0.0622) |
| 2012 | -0.6249 | (-0.7338, -0.5156) | 12.5240 | (5.1599, 22.6408) | 0.0404 | (0.0218, 0.0562) |
| 2013 | -0.5035 | (-0.5995, -0.4095) | 12.7588 | (5.5134, 22.7535) | 0.0398 | (0.0203, 0.0547) |
| 2014 | -0.8214 | (-0.9197, -0.7222) | 12.8010 | (5.5933, 22.8088) | 0.0424 | (0.0242, 0.0604) |
| 2015 | -0.7063 | (-0.7908, -0.6185) | 12.7208 | (5.8189, 22.6522) | 0.0458 | (0.0309, 0.0736) |

**Table 4.** Point estimates and 95% CIs for mean and proportion parameters for *Salmonella* I,4,[5],12:i:- tested with (TIO) in the ISU VDL dataset.

|  | Estimated mean log<sub>2</sub>MIC in the non-resistant population |  | Estimated mean log<sub>2</sub>MIC in the resistant population |  | Estimated proportion of resistant population |  |
| --- | --- | --- | --- | --- | --- | --- |
| Year | $\hat{\beta}_{0i}$ | 95% CI of $\hat{\beta}_{0i}$ | $\hat{\beta}_{1i}$ | 95% CI of $\hat{\beta}_{1i}$ | $\hat{p}_i$ | 95% CI of $\hat{p}_i$ |
| 2011 | -1.5938 | (-2.0234, -1.1577) | 3.0631 | (2.6063, 3.5410) | 0.2214 | (0.1648, 0.2733) |
| 2012 | -1.4962 | (-1.6860, -1.3145) | 3.3031 | (3.0435, 3.6241) | 0.2126 | (0.1574, 0.2536) |
| 2013 | -1.0193 | (-1.1410, -0.8924) | 3.3055 | (3.0956, 3.5459) | 0.2100 | (0.1645, 0.2472) |
| 2014 | -0.9867 | (-1.1124, -0.8591) | 3.2542 | (3.0673, 3.4813) | 0.2100 | (0.1627, 0.2428) |
| 2015 | -0.8984 | (-1.0210, -0.7761) | 3.0831 | (2.9517, 3.2393) | 0.2341 | (0.2011, 0.2842) |
| 2016 | -0.9169 | (-0.9952, -0.8391) | 3.1126 | (3.0195, 3.2254) | 0.2288 | (0.2043, 0.2628) |
| 2017 | -0.6747 | (-0.7633, -0.5882) | 3.0487 | (2.9413, 3.1770) | 0.2137 | (0.1804, 0.2411) |

**Table 5.** Point estimates and 95% CIs for linear model parameters in the three examples.

|  |  | Estimated intercept for the linear trend of non-resistant mean log<sub>2</sub>MIC |  | Estimated slope for the linear trend of non-resistant mean log<sub>2</sub>MIC |  |
| --- | --- | --- | --- | --- | --- |
| Data set | Antibiotic | $\hat{\gamma}_0$ | 95% CI of $\hat{\gamma}_0$ | $\hat{\gamma}_1$ | 95% CI of $\hat{\gamma}_1$ |
| CDC NARMS | CHL | 1.6225 | (1.3416, 1.8861) | 0.0415 | (0.0088, 0.0773) |
| CDC NARMS | TIO | -1.2297 | (-1.6223, -0.7820) | 0.0369 | (-0.0168, 0.0844) |
| ISU VDL | TIO | -1.6391 | (-2.2577, -1.0546) | 0.1388 | (0.0049, 0.2713) |

**Fig 3.**
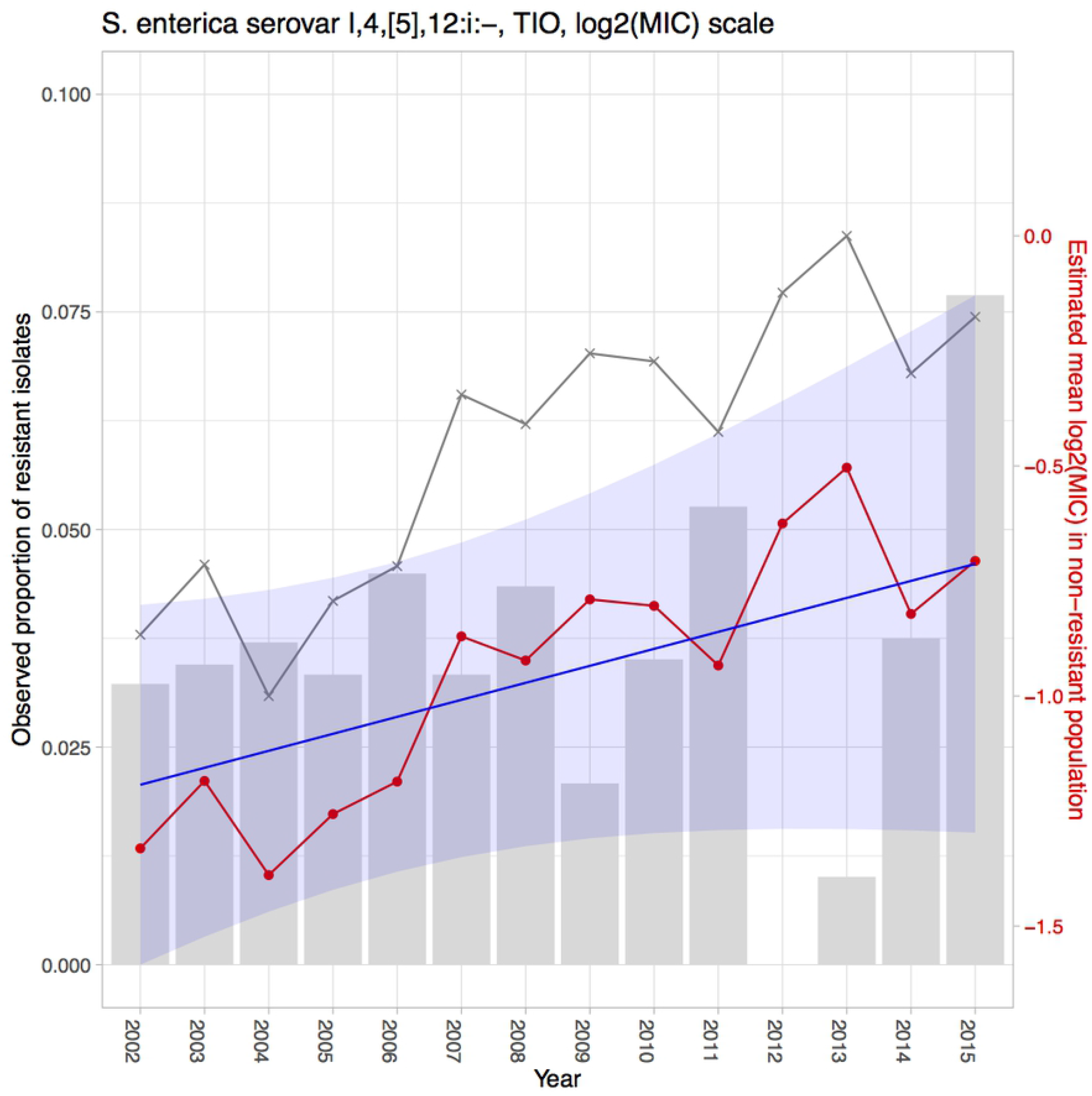
Estimation results for *Salmonella* enterica I,4,[5],12:i:- tested with TIO in the CDC NARMS dataset. The grey bars represent the observed proportions of resistant bacteria (left y-axis). The grey line indicates the naïve mean of log_2_MIC in the non-resistant population. The red line connects the point estimates of the mean log_2_MIC in the non-resistant population. The blue line represents the estimated linear trend, shaded with its 95% CI.

**Fig 4.**
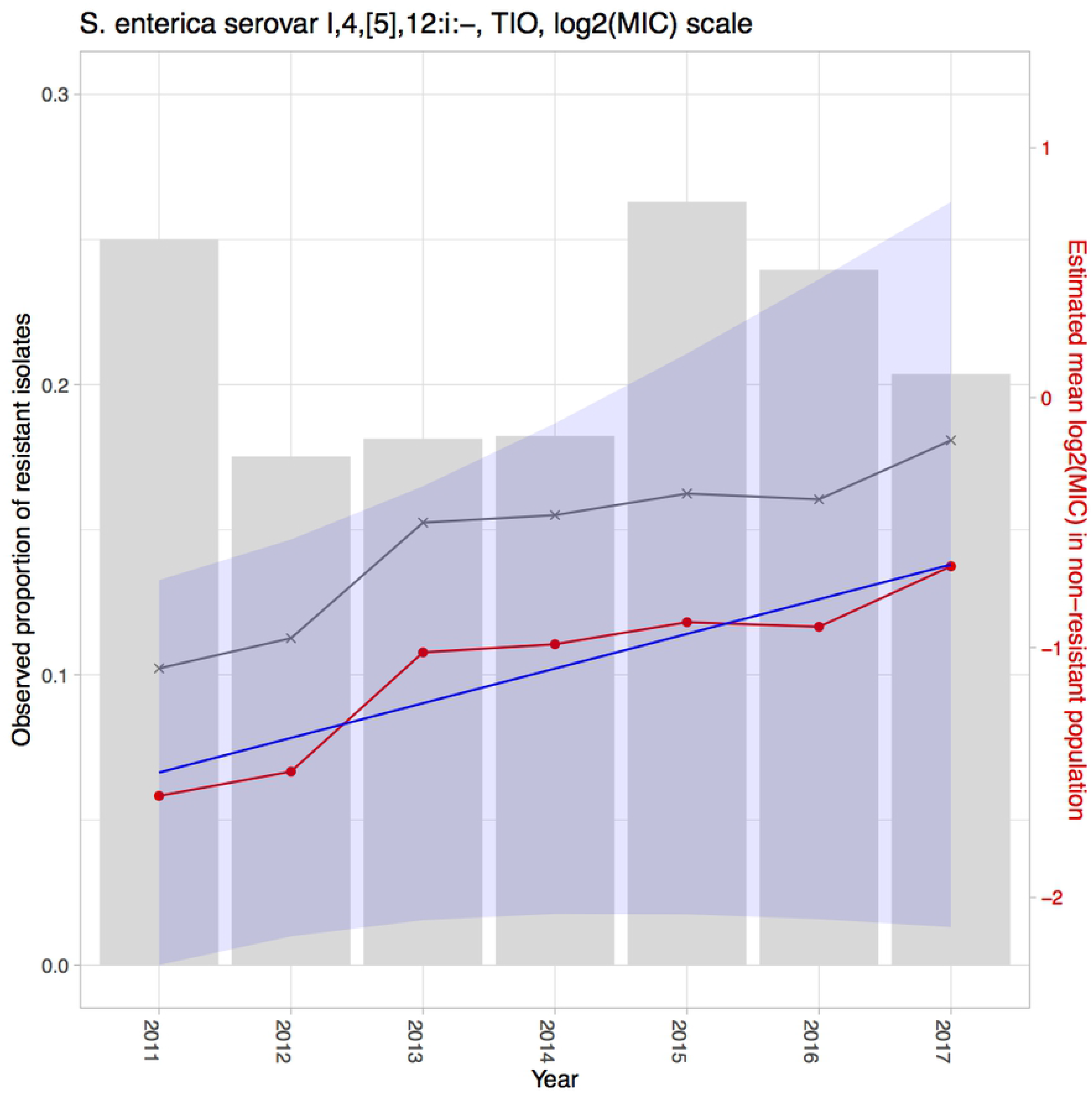
Estimation results for *Salmonella* enterica I,4,[5],12:i:- tested with TIO in the ISU VDL dataset. The grey bars represent the observed proportions of resistant bacteria (left y-axis). The grey line indicates the naïve mean of log_2_MIC in the non-resistant population. The red line connects the point estimates of the mean log_2_MIC in the non-resistant population. The blue line represents the estimated linear trend, shaded with its 95% CI.

## Results

Using the proposed hierarchical Bayesian latent class mixture model with censoring and linear trend, we analyzed the resistance level of *Salmonella* enterica I,4,[5],12:i:- tested with CHL and TIO in the CDC NARMS human data and also *Salmonella* enterica I,4,[5],12:i:- tested with TIO in the ISU VDL swine dataset. The goal of the analyses was to illustrate the evaluation of increasing or decreasing trends in mean log_2_MIC over time in order to identify trends with important public health implications. The estimates of the yearly mean log_2_MIC in both the non-resistant 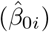 and resistant populations 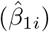 and estimates of the proportions of the population designated as resistant 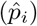, along with their 95% CIs, are presented in Table 2, Table 3, and Table 4. A linear model was fitted to the mean log_2_MIC in the non-resistant population to borrow information across years and to reveal a potential linear trend. The intercept (*γ*_0_) was interpreted as the baseline of the mean log_2_MIC in the non-resistant population, while the slope (*γ*_1_) was interpreted as an increase in the resistance from the previous year (for *i* = 2, …, *I*) or from the baseline (for *i* = 1). The point and interval estimates of the linear model parameters for each example are presented in Table 5. Among all the above-mentioned estimates, the estimates for the yearly mean log_2_MIC 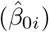 and the linear parameters 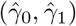 of the non-resistant population are of the greatest interest, as the main objective of our study was to use the proposed model to detect linear temporal changes in AMR in the susceptible group. If 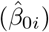 shows an increase over time and the estimated slope 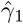 is positive, then this result could signify increasing resistance for the organism to the antibiotic. The estimates for these parameters are also presented in Fig 2, Fig 3, and Fig 4 for better visualization. The yearly mean log_2_MIC in the resistant population 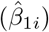 was not our priority in this study, as most of the observations in the resistant population are right censored and thus do not provide enough information for parameter estimations. This right censoring also underlay the wide credible intervals of 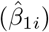.

From the first row in Table 5, which shows the estimations of the linear model parameters for *Salmonella* I,4,[5],12:i:- tested with CHL in the CDC NARMS dataset, the 95% CI for the slope was (0.0088, 0.0773), indicating a significantly positive slope estimation and therefore a significantly increasing trend in the mean log_2_MIC of the non-resistant population. Fig 2 depicts this increasing pattern by plotting the fitted regression line (blue line) through the estimated non-resistant means (red points). The yearly non-resistant means were scattered around the regression line in a random pattern, in agreement with the linear model assumption of independence. The grey histogram in Fig 2 shows the observed proportions of the resistant isolates. Notably, no *Salmonella* I,4,[5],12:i:- isolate resistant to CHL was observed for some years, while the mean of log_2_MIC below the resistance threshold was estimated to be increasing constantly. This example demonstrated a situation in which analysis based on dichotomized MIC alone would misleadingly indicate a decreasing level of resistance from 2009 to 2012 and neglect the increasing level of resistance below the break point. Based on these results, we concluded that an intervention for use of CHL for *Salmonella* I,4,[5],12:i:- in human is suggested to prevent a possible outbreak of resistance if the linearly increasing pattern is allowed to continue in the following years.

In the example of *Salmonella* I,4,[5],12:i:- tested with TIO in the CDC NARMS dataset, we found an insignificant slope estimation, with a 95% CI of (−0.0168, 0.0844) (second row in Table 5). Despite the notion the true value of the slope parameter is within an interval that contains zero, our best estimation was positive, and the major coverage of the CI was greater than zero. No organism exhibited resistance to TIO above the threshold in 2012, reflecting a rapid decrease from more than 5% in 2011. This phenomenon was accompanied by a stable MIC increase in the non-resistant population.

For the ISU VDL dataset, we detected a significantly increasing pattern in the non-resistant means for *Salmonella* I,4,[5],12:i:- tested with TIO, with a 95% CI of (0.0049, 0.2713) (third row in Table 5). As shown in Fig 4, the 95% CI of the estimated regression line (the shaded area) is rather wide compared with those in Fig 2 and Fig 3, and the difference likely resulted from the limited number of observations in this example. In this VDL swine dataset, the observed resistant proportions exhibited an abrupt decrease in 2012, an abrupt increase in 2015, and a continuous decrease since that time. Our estimated regression line added more dimensions in the non-resistant population by revealing the creep in its mean log_2_MIC.

## Model evaluation with simulation

In order to assess the performance of the proposed hierarchical Bayesian latent class mixture model with censoring and linear trend, a simulation study was conducted based on the CDC NARMS-CHL example. In our model, the parameters of interest were 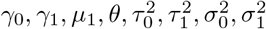, and *ν*^2^. In the simulation study, these parameters were pre-determined according to the estimation results from the example and were denoted with a “hat” on the Greek letter. Also, we simulated the same number of observations for the *i*th year as the total number of isolates (*n_i_*) in the CDC NARMS-CHL dataset. The data generation process is described in the following steps.

For *i* = 1, 2, …, *I*; *j* = 1, 2, …, *n*_*i*_; and *t*_*i*_ = *i*:

1. Generate 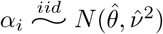.
2. Calculate *p*_*i*_ by performing an expit transformation (inverse of logit) on *α*_*i*_, 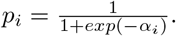.
3. Generate the latent class indicator 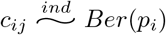.
4. Generate the mean log_2_MIC in the non-resistant population 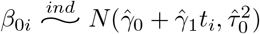.
5. Generate the mean log_2_MIC in the resistant population 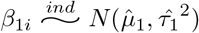.
6. Generate the log_2_MIC for isolate *j* in year *i* with 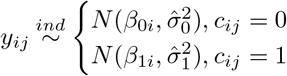.
The simulated *p*_*i*_ and *β*_0*i*_ were treated as the real proportion of resistant population and the real yearly mean of log_2_MIC, which were compared with estimations based on the simulated observations subsequently. Until this point, the simulated *y*_*ij*_ were continuous values drawn from a mixture of Gaussian distributions. To mimic the censoring nature of log_2_MIC, *y*_*ij*_ was also censored according to its value. We previously defined *l*_*ij*_ and *u*_*ij*_ to be the lower bound and upper bound of *y*_*ij*_. For CHL, the starting dilution was 2 mg/ml, and the ending dilution was 32 mg/ml for both serotypes. This indicated that if *y*_*ij*_ ≤ log_2_(2), then it will be left censored as *y*_*ij*_ ≤ 1 with *l*_*ij*_ = −∞ and *u*_*ij*_ = 1. Similarly, if *y*_*ij*_ > log_2_(32), it will be right censored as *y*_*ij*_ > 5, with *l*_*ij*_ = 5 and *u*_*ij*_ = ∞. If log_2_(2) < *y*_*ij*_ ≤ log_2_(32), then *y*_*ij*_ will be interval censored with *l*_*ij*_ as its nearest integer to the left and *u*_*ij*_ as its nearest integer to the right. This censoring operation corresponds to step 7.
7. Censor the underlying true values of log_2_MIC, *y*_*ij*_, to the observed values 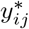 with 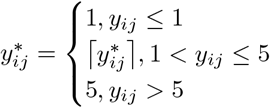, where 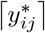 represents rounding up to the nearest integer. End of simulation.

As a single set of observed log_2_MIC (i.e., 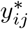) could be generated by completing steps 1 to 7, we simulated 100 datasets by repeating the above procedure 100 times. With each simulated dataset, estimation was conducted using the proposed hierarchical Bayesian model, which produced a set of estimations: 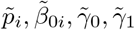, etc., for *i* = 1, 2, …, *I*. In order to assess the performance of our hierarchical Bayesian model, two metrics were calculated. The first was the mean bias over the 100 simulations, and the second was the root of mean squared error (RMSE). Mean bias measures how close the estimations are relative to the true parameter values, while RMSE measures the variation of estimates around the true parameters. Table 6 shows the mean bias and RMSE for the yearly parameters, and Table 7 shows the mean bias and RMSE for the linear parameters. The mean biases and RMSE for *p_i_*, *β*_0*i*_, *γ*_0_, and *γ*_1_ were very close to 0 compared with the magnitude of their own estimations, indicating precise and robust estimations for the most relevant parameters utilizing our proposed model.

**Table 6.**
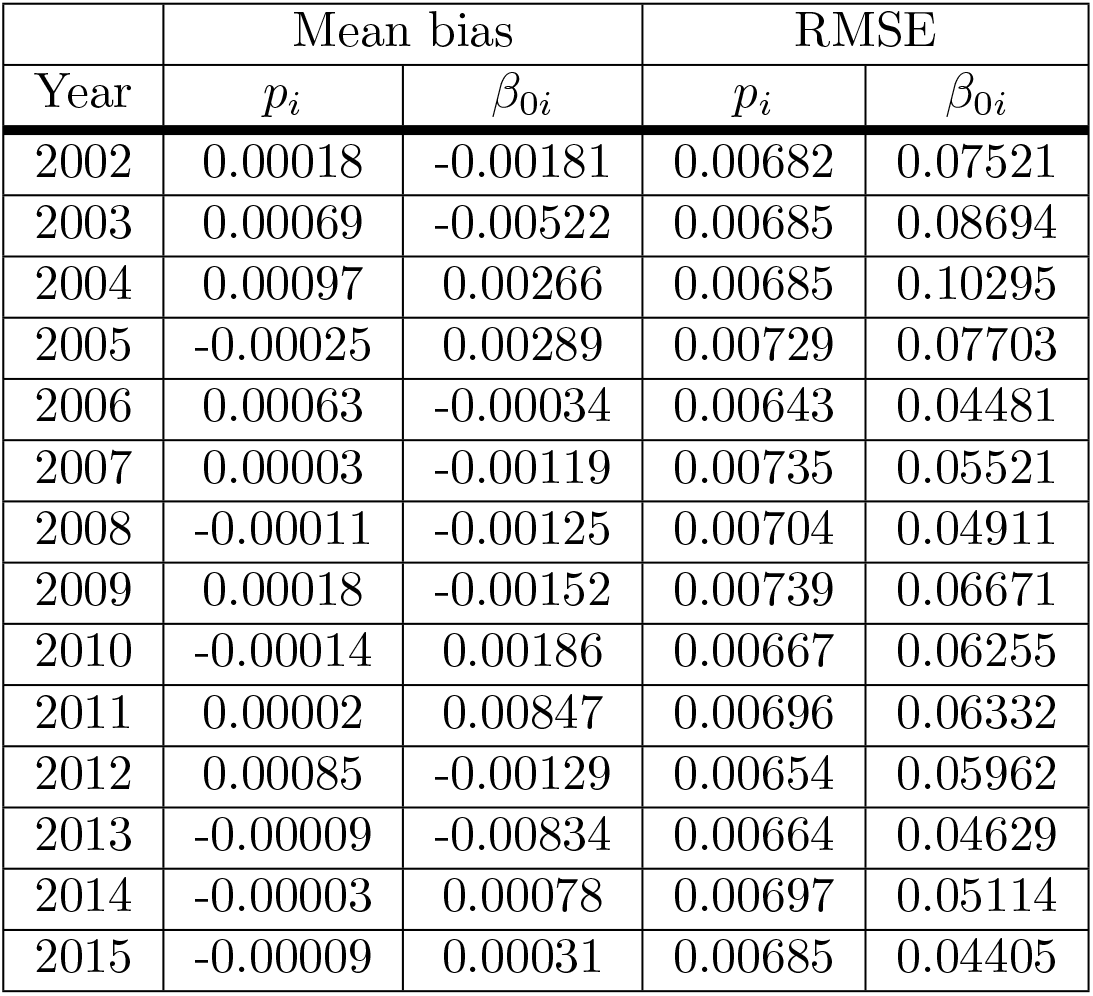
The mean bias and RMSE of the proportion and mean parameters from the simulation study.

| Year | Mean bias |  | RMSE |  |
| --- | --- | --- | --- | --- |
| | $p_i$ | $\beta_{0i}$ | $p_i$ | $\beta_{0i}$ |
| 2002 | 0.00018 | -0.00181 | 0.00682 | 0.07521 |
| 2003 | 0.00069 | -0.00522 | 0.00685 | 0.08694 |
| 2004 | 0.00097 | 0.00266 | 0.00685 | 0.10295 |
| 2005 | -0.00025 | 0.00289 | 0.00729 | 0.07703 |
| 2006 | 0.00063 | -0.00034 | 0.00643 | 0.04481 |
| 2007 | 0.00003 | -0.00119 | 0.00735 | 0.05521 |
| 2008 | -0.00011 | -0.00125 | 0.00704 | 0.04911 |
| 2009 | 0.00018 | -0.00152 | 0.00739 | 0.06671 |
| 2010 | -0.00014 | 0.00186 | 0.00667 | 0.06255 |
| 2011 | 0.00002 | 0.00847 | 0.00696 | 0.06332 |
| 2012 | 0.00085 | -0.00129 | 0.00654 | 0.05962 |
| 2013 | -0.00009 | -0.00834 | 0.00664 | 0.04629 |
| 2014 | -0.00003 | 0.00078 | 0.00697 | 0.05114 |
| 2015 | -0.00009 | 0.00031 | 0.00685 | 0.04405 |

**Table 7.** The mean bias and RMSE of the linear model parameters from the simulation study.

| Mean bias |  | RMSE |  |
| --- | --- | --- | --- |
| $\gamma_0$ | $\gamma_1$ | $\gamma_0$ | $\gamma_1$ |
| -0.02105 | 0.00330 | 0.20174 | 0.02669 |

## Discussion

Our goal with this study was to address a deficiency in the available approaches for analysis of AMR data using MIC values. AMR is a serious public health issue worldwide, and enormous resources have been devoted to monitoring the changes in MIC over the years. We proposed a Bayesian hierarchical model with a linear trend and demonstrated that this model enables additional information about the linear patterns in the mean log_2_MIC in the non-resistant population in addition to the proportion of resistant bacteria. The linear changes in the non-resistant population may be linear creep that signals a need for intervention or a linear decline that implies a successful intervention. Therefore, our proposed model offers a tool to a more complete picture of the resistance level of organisms against various antibiotics, providing valuable information for the surveillance programs.

The proposed approach was founded on the concept of identifying a valid and robust mean log_2_MIC estimate that addresses the latent nature of the MIC data. Under our framework, there were two sources of latent parameters. One resulted from censorship, as the true underlying continuous values for the censored observed MIC values are unknown. The other involved the population from which the bacterial isolates arose (resistant or non-resistant population). By tackling the censorship problem and incorporating the mixed components of the data, our Bayesian hierarchical model corrected the systematic bias of the mean MIC estimations and separated the isolates from different groups. We then added a higher level of complexity to this fundamental model setup: linear regression in the hierarchical model.

Considering the variation of the MIC values shown in the dataset, we allowed the mean log_2_MIC to vary across different years. The Bayesian hierarchical model yielded a more robust estimation by shrinking the mean estimates towards a linear regression line in the non-resistant population and towards a common constant in the resistant population. In this way, we were able to quantify the linear pattern in the mean log_2_MIC in the group of isolates that are often undervalued by researchers. Compared with regressing the mean log_2_MIC to a constant in the non-resistant population, a linear trend with a non-zero slope provided a better fit to the datasets and satisfied our model assumptions.

Our model relied on several assumptions. We assumed normal distributions for resistant and non-resistant populations. This assumption was supported by the observed MIC distribution for the examples used in this paper; however, under violation of this assumption, non-parametric methods, such as spline fitting, could be used to replace the normality assumption [1]. For both resistant and non-resistant populations, we also assumed invariant variances across years, following the principle of parsimonious models. This assumption could be important when there are observations from many years but not enough observations within each year. In addition, we assumed that the proportion of the resistant population was independent across all years. We also assumed the mean log_2_MIC had independent errors in the linear model and the constant mean model in the sub-populations. Violation of this assumption would require inclusion of a correlation structure in the proportions or the errors terms.

In conclusion, we proposed a framework of analysis of longitudinal log_2_MIC data using a Bayesian hierarchical approach with linear trend. We not only estimated the mean of log_2_MIC values properly and accurately but also detected a significant linear increase in the mean log_2_MIC in the non-resistant population for some given organisms and antibiotics, potentially signaling the need for intervention. Additional directions from this proposed framework include studying the correlations among multiple antibiotics, between human and animal resistance, and between different surveillance programs for the same population. In addition, analysis of the relationship between clinical interventions and the MIC responses to the interventions based on these models is of interest.

## Acknowledgments

We are grateful to Dr. Chaohui Yuan for providing help with the R scripts for the basic model without the linear temporal trend.

## Supporting information

### S1 Appendix. Gibbs sampling procedure

The Gibbs sampling procedure was conducted as described here. We used “· | ·” to denote full conditional distribution unless otherwise specified. This parameter is the distribution of what is before the pipe and is conditional on all other parameters involved in the model. In the following steps, *j* = 1, …, *n*_*i*_, and *i* = 1, …, *I*.

1. Obtain draws of latent continuous variable *y*_*ij*_ from censored observation 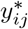 using the inverse cumulative distribution function (inverse CDF) method. By observing 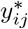 we sampled from the full conditional normal distribution with the boundaries *l*_*ij*_ and *u*_*ij*_. To be more specific, the three censoring situations are discussed below:

- When 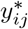 is interval censored with limits *l*_*ij*_ and *u*_*ij*_, *y*_*ij*_ is updated via

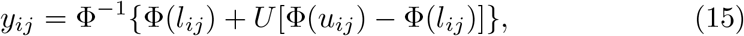

where Φ is the CDF function of standard normal distribution, and Φ^−1^ is the inverse of the CDF function. *U* is a random draw from *Unif*(0, 1).
- When 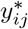 is left censored with limits *l*_*ij*_ = −∞ and *u*_*ij*_, *y*_*ij*_ is updated via

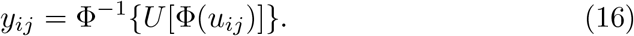
- When 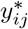 is right censored with limits *l*_*ij*_ and *u*_*ij*_ = ∞, *y*_*ij*_ is updated via

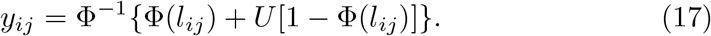
2. Draw samples of *c*_*ij*_ from their full conditional distribution

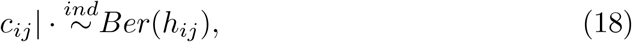

where 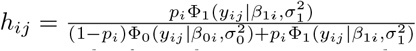 describing the chance for an observation to be from the resistant population. *ϕ*(*y*|*β, σ*^2^) represents the probability density function of a normal distribution with mean *β* and variance *σ*^2^.
3. Sample the intercept parameter *γ*_0_ in the linear part from its full conditional distribution

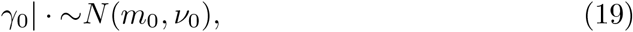

where 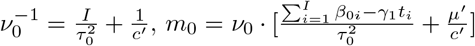, *μ*′ = 0 is the prior mean of *γ*_0_, and *c*′ = 10^6^ is the prior variance of *γ*_0_.
4. Sample the slope parameter *γ*_1_ in the linear part from its full conditional distribution

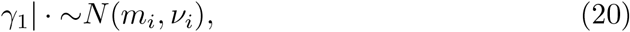

where 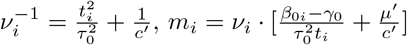, *μ*′ = 0 is the prior mean of *γ*_1_, and *c*′ = 10^6^ is the prior variance of *γ*_1_. Starting from *i* = 1, for *i* < *I*, return to step 3 and 4 to sample linear parameters for another time. Increase *i* by 1. When *i* reaches *I*, continue to step 5.
5. Sample *μ*_1_, the hierarchical yearly mean of the mean log_2_MIC in the resistant population, from its full conditional distribution

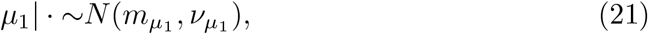

where 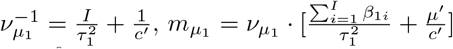 *μ*′ = 0 is the prior mean of *μ*_1_, and *c*′ = 10^6^ is the prior variance of *μ*_1_.
6. Sample the real yearly mean log_2_MIC in the non-resistant population, *β*_0*i*_, from its full conditional distribution

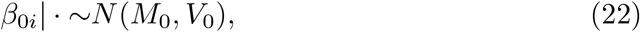

where 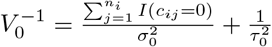 and 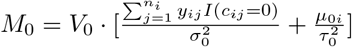
7. Sample the real yearly mean log_2_MIC in the resistant population, *β*_1*i*_, from its full conditional distribution

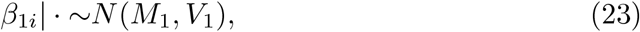

where 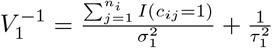 and 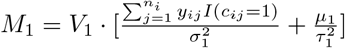
8. Sample 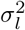 variance of latent log_2_MIC, in either population from its full conditional distribution

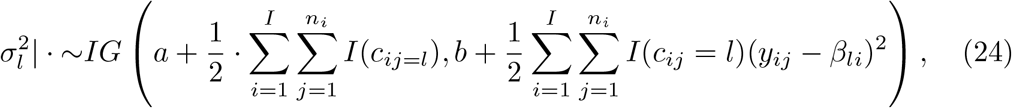

where *l* = 0 represents the non-resistant population, and *l* = 1 represents the resistant population. *a* = 10^−4^ is the prior shape parameter, and *b* = 10^−4^ is the prior rate parameter.
9. Sample 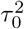 the variance for the mean log_2_MIC in the non-resistant population, from its full conditional distribution

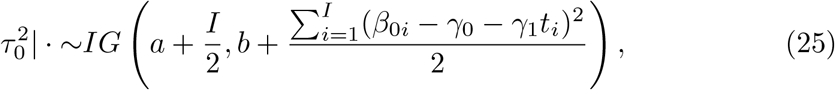

where *a* = 10^−4^ is the prior shape parameter, and *b* = 10^−4^ is the prior rate parameter.
10. Sample 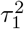 the variance for the mean log_2_MIC in the resistant population, from its full conditional distribution

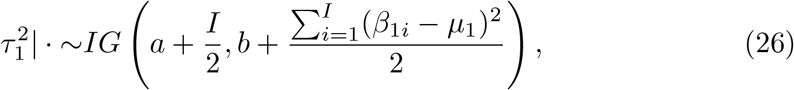

where *a* = 10^−4^ is the prior shape parameter, and *b* = 10^−4^ is the prior rate parameter.
11. Obtain draws of parameter *θ* from its full conditional distribution

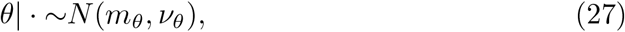

where 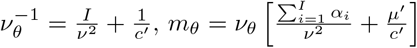 *μ*′ = 0 is the prior mean of *θ*, and *c*′ = 10^6^ is the prior variance of *θ*.
12. Sample *ν*^2^, the variance of *α_i_*, from its full conditional distribution

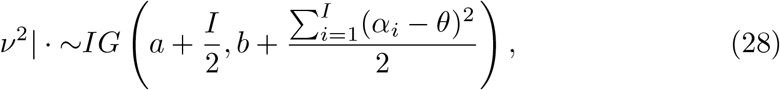

where *a* = 10^−4^ is the prior shape parameter, and *b* = 10^−4^ is the prior rate parameter.
13. Sample *α_i_*, the log-odds of the proportion, using random walk Metropolis-Hastings from their posterior to some constant

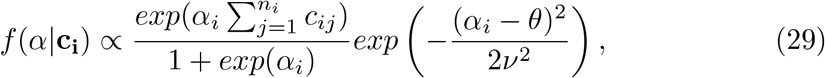

where **c**_**i**_ is the vector of *c*_*ij*_, *j* = 1, …, *n_i_*.

### S1 File. Editing certificate

The certificate of English editing is attached in the Supporting information as an external file.

## Footnotes

1 Github repository: https://github.com/MinZhang95/AMR-Linear

